# A Pilot Study Investigating the Impact of Oral Probiotic Supplementation on the Vaginal Microbiota and Aspects of Well-being

**DOI:** 10.64898/2026.10.24.760943

**Authors:** Eleri Hulme, Daniel A. John, Daryn R. Michael, Niall Coates, Joshua M.D. Smith, Elwira Sośniak, Esther Gimblett, Maya Dabcheva, Duolao Wang, Sue F. Plummer

## Abstract

This double-blind, randomised, placebo-controlled pilot study investigated the effects of a six-strain probiotic formulation containing *Lactobacillus crispatus* CUL40 and *L. gasseri* CUL09 on vaginal microbiota composition and well-being in 40 women with recurrent vaginal discomfort. Participants received either the probiotic (20 billion colony-forming units/day) or placebo for 20 days. Vaginal samples collected at baseline, midpoint and endpoint were analysed using shotgun metagenomic sequencing, microbial culture and species- and strain-specific quantitative polymerase chain reaction. Vaginal symptoms and broader well-being were recorded daily. At endpoint, vaginal detection of CUL40 and CUL09 was significantly higher in the probiotic group, with neither strain detected in the placebo group. Viable *L. crispatus* CUL40 was recovered from vaginal cultures, whereas viable CUL09 was not recovered. *L. gasseri* relative abundance was significantly higher in the probiotic group at midpoint and endpoint, while Gardnerella counts were significantly lower at endpoint. Probiotic supplementation significantly influenced beta diversity over time between groups, although community state types remained broadly stable. During the second half of the intervention, rates of vaginal itchiness, burning sensation and anxiousness were significantly lower in the probiotic group. Bloating was also less frequent throughout the intervention. These findings demonstrate increased vaginal detection of orally administered probiotic strains, including recovery of viable CUL40, alongside changes in vaginal microbiota composition and reduced vaginal discomfort. Larger studies are warranted to confirm these exploratory findings and establish longer-term effects. This trial was registered at ClinicalTrials.gov (NCT07364383).

**Importance:** Evidence that probiotic bacteria taken orally can subsequently be detected alive in the vagina remains limited. In this pilot trial, we combined genetic testing with bacterial culture to investigate vaginal detection of *Lactobacillus crispatus* CUL40 and *Lactobacillus gasseri* CUL09 following oral supplementation. Both strains were detected, and viable CUL40 bacteria were recovered. These findings were accompanied by changes in vaginal bacterial communities and reduced reporting of some symptoms. The study provides evidence to guide further research into oral probiotics and vaginal health.

## Introduction

The vaginal tract is a hormonally responsive mucosal environment that harbours a locally protective, taxonomically conserved microbial community that tends to be rich in *Lactobacillus* spp. The vaginal microbiota plays an important role in creating an acidic environment to support colonisation resistance and exclude potential pathogens from the vaginal ecosystem(1–3). The microbiota interacts continuously with the vaginal epithelium, mucus and local immune system to create a balanced environment and five main classes of microbiota composition have been identified. These classes are known as Community State Types (CSTs) that are classified according to the dominant organism present; CST I: *Lactobacillus crispatus*, CST II: *L. gasseri,* CST III: *L. iners,* CST V: *L. jensenii,* whereas CST IV encompasses more diverse, *Lactobacillus*-depleted communities, often enriched in anaerobic bacteria and is often considered to be a less favourable category(4, 5).

The stability of the vaginal microbiota is subject to challenges from menstruation and the associated hormonal and pH fluctuations, antibiotic use, excessive hygiene practices and sexual activity(2, 6). The microbiota composition does not change significantly in response to hormonal challenges alone but the cumulative effects of challenges can disrupt the balance and open the path for potential pathogens. Those with the more diverse CST IV population are considered to be more at risk of challenge and, although many women are asymptomatic, CST IV is often associated with bacterial vaginosis (BV)(7–9).

Probiotics (live microorganisms that, when administered in adequate amounts, confer a health benefit on the host(10)) have been widely used to support gut health but are becoming recognised for their beneficial impacts throughout the body. Several clinical studies have investigated the oral administration of *Lactobacillus* strains, with reported benefits including changes in vaginal microbiota composition, reduced recurrence of BV and alleviation of vaginal symptoms(11–14). Orally administered bacteria are thought to reach the vaginal environment following gastrointestinal transit, with subsequent transfer from the intestinal or rectal reservoir to the vagina via the perineal region (vaginal-rectal axis)(15–17). There are differences in the *Lactobacillus* species and strains typically found in the vaginal and gastrointestinal tracts and it is important to determine whether orally administered, vaginally-derived, probiotic strains can reach and become established in the vaginal tract.

The Lab4 probiotic consortium has been found to exert beneficial effects in healthy women but its impact within the vaginal ecosystem has not been assessed to date(18–20). The Lab4V consortium is a tailored version of Lab4 intended to enhance its suitability to support the vaginal microbiota by the addition of *Lactobacillus crispatus* CUL40 and *Lactobacillus gasseri* CUL09 and this study observed the effects of oral supplementation of the formulation in women with a history of recurrent vaginal discomfort. Changes in vaginal microbiota composition were assessed, including the translocation of the supplemented vaginal strains, together with changes in vaginal symptom incidence and in markers of both intestinal health and broader aspects of wellbeing.

## Methods

### Study Design

This was a double-blind, randomised, placebo-controlled pilot study in which 40 participants were allocated equally to two parallel groups.

### Study Approval

The study was performed in accordance with the principles of the Declaration of Helsinki. Ethical approval was granted by the Ethical Committee of Clarenta (Reference: No.001/27.11.2025r). The study was prospectively registered with ClinicalTrials.gov: NCT07364383.

### Recruitment & Randomisation

The inclusion criteria; healthy females (at birth), 18 to 40 years old with a history of 3 or 4 episodes of vaginal discomfort (1 or more days) involving itching, burning sensation and/or odorous discharge in the previous 12 months. Willing to start on completion of monthly cycle (< 5 days post bleeding), provide vaginal and faecal samples as required, maintain normal diet and lifestyle, refrain from taking other probiotic supplements during the study and, for the 2 days prior to providing vaginal swabs, refrain from sexual activity, the use of lubricants and vaginal cleaning (with antibacterials)/douching. The exclusion criteria included any oral antibiotic intake during previous 2 months; pregnant or planning pregnancy; given birth in the last 3 months; experiencing a vaginal or urinary infection; suffering with immunodeficiency/receiving immunosuppressive therapy or diagnosed with diabetes/cardiovascular disease/cancer/dementia.

### Study Product

The probiotic capsules comprised the Lab4V consortium of *Lactobacillus acidophilus* CUL60, *L. acidophilus* CUL21, *L. crispatus* CUL40, *L. gasseri* CUL09, *Bifidobacterium bifidum* CUL20 and *B. animalis* subsp. *lactis* CUL34 (20 billion colony-forming units (CFU)), vitamin A (800 IU) and vitamin D3 (1000 IU) on a microcrystalline cellulose base; the matched placebo capsules contained microcrystalline cellulose (Cultech Ltd, Port Talbot, UK). Participants took one capsule daily (with or following food but not with hot drinks) for 20 days, stored capsules in a refrigerator and returned unused capsules for compliance monitoring.

### Study aims

The objectives of this short-term exploratory study were to assess the impact of probiotic supplementation on the composition of the vaginal microbiota, including whether the vaginal probiotic strains could be detected within the vaginal ecosystem, and to monitor participant wellbeing.

### Data & Sample collection

The sequence of data and sample collection during the study are shown in Figure 1A. Anthropometric measurements were taken at baseline. Vaginal swabs (and stool samples) were collected at baseline, at midpoint (Day 10) and endpoint (Day 20). Wellbeing diaries were completed daily.

**Figure 1.**
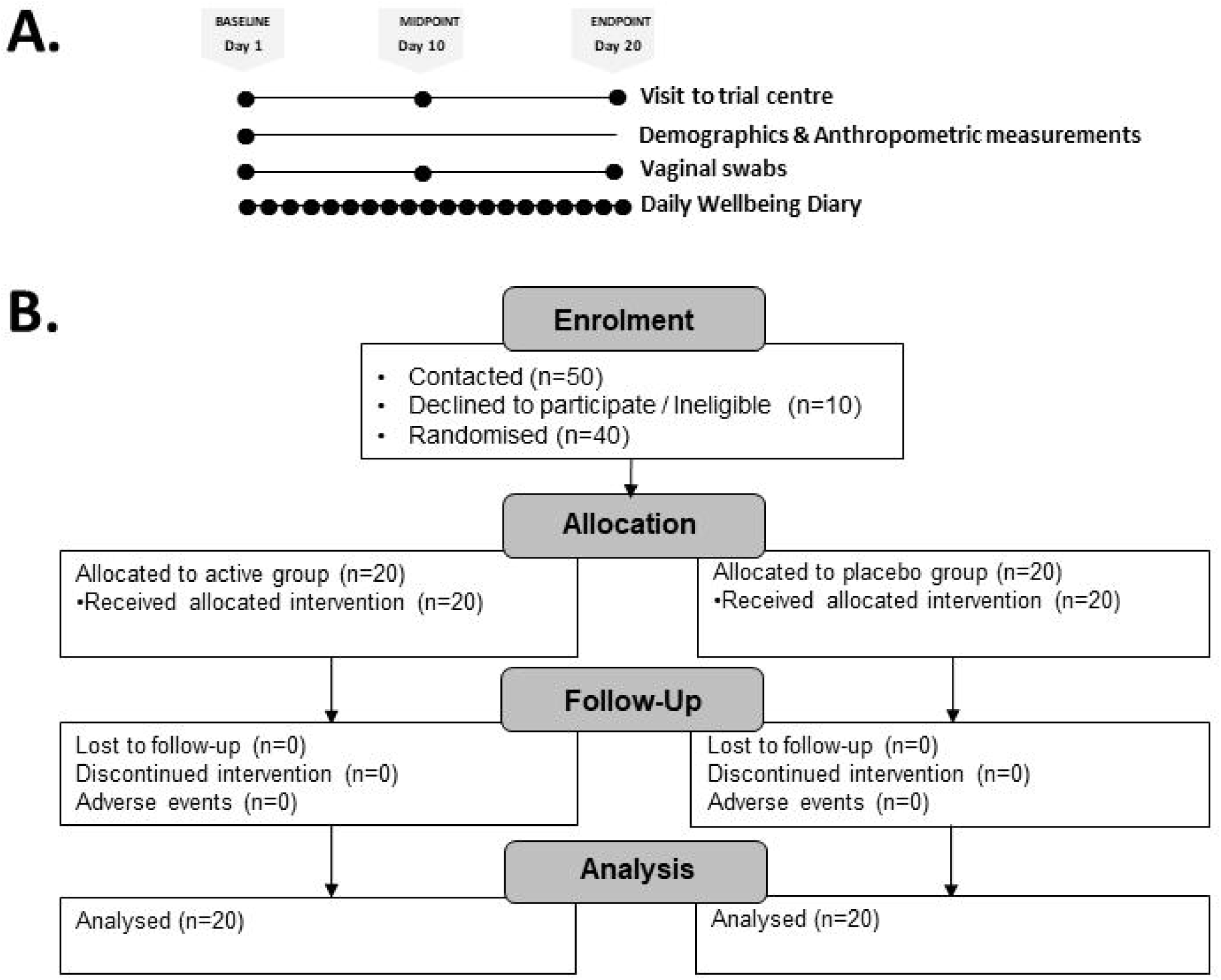
Overview of the Study. (A) Sequence data and sample collection and (B) a study flow diagram

### Weight/Height measurement

Body weight was measured after the removal of shoes and jackets using a calibrated column scale (Seca 709, Hamburg, Germany). Shoes were removed before height measurement.

### Vaginal swabbing

Vaginal samples were collected from participants who were not actively menstruating using sterile swabs (Thermo Scientific Swab Plastic PlainTube F155Ca). Samples were frozen at -80°C within 5 minutes of collection. For processing, the frozen swab heads were transferred into 1 mL of Maximum Recovery Diluent (MRD, ThermoFisher), vortexed for 20s, incubated at 5°C for 10 min, and the vortex–incubation step was repeated twice. 150µL of the suspension was used for microbial culture on selective media and 250µL was used for genomic analysis. The remaining sample suspension was stored at −80°C.

### Genomic DNA Extraction from vaginal swabs

The ZymoBIOMICS Spike-in Control, comprising *Imtechella halotolerans* and *Allobacillus halotolerans* (Zymo Research, United States), was added to each sample prior to DNA extraction as an internal standard for shotgun metagenomic sequencing. Genomic DNA (gDNA) was extracted from vaginal samples using the QIAamp® PowerFecal Pro DNA Kit (Qiagen, Germany). Eluted gDNA was stored at −80°C until analysis.

### Shotgun Metagenomic Sequencing

Shotgun metagenomic sequencing of the vaginal gDNA extracts was performed using the Illumina NovaSeq X platform (Novogene, UK) with 150-nucleotide paired-end reads. Quality control and host read removal of raw sequences were performed using KneadData (v0.12.2)(21). The resulting host-depleted, quality-filtered reads were retained for downstream metagenomic analyses. Taxonomic profiles were generated using Kraken2 (v.2.1.3)(22) against the Vaginal VMGC custom database v1(23), with the addition of the T2T CHM13 human genome, the ZymoBIOMICS spike-in bacteria and complete PacBio assemblies of the supplemented probiotic organisms. Bracken (v.3.0.1)(24) was used to refine species-level abundance estimates. Species-level read counts were normalised to the number of *Imtechella halotolerans* spike-in reads detected in each sample, after which spike-in organisms were removed from the dataset prior to downstream microbiota analyses. CSTs were assigned using VALENCIA(4); CST I (*Lactobacillus crispatus*-dominated), CST II (*L. gasseri*-dominated), CST III (*L. iners*-dominated), CST IV (heterogeneous, Lactobacillus-depleted communities) and CST V (*L. jensenii*-dominated). Shannon and Simpson alpha diversity indices were calculated using phyloseq (v1.50.0(25)) before abundance and prevalence filtering. Bacterial species with at least 0.01% relative abundance in 15 or more samples for beta diversity and differential abundance/prevalence analyses.

### Microbial culture of vaginal swabs

Viable counts were determined by culture on selective media (Supplementary Table S1). Bacterial counts were expressed as log colony-forming units (CFU)/swab. The MRS plates were retained for colony PCR analysis.

### Detection of *L. crispatus* and *L. gasseri* species and the *L. crispatus* CUL40 and *L. gasseri* CUL09 strains in vaginal samples by PCR

Design of strain-specific primers for CUL40 (Forward: 5’-TGGACTGATTCCCCCTTTGAC-3’, Reverse: 5’ TGGGAGGTCATCAACAACGTAG-3’) and CUL09 (Forward: 5’-TATGGTCGGTTAACACGGGTC-3’, Reverse: 5’-ACGGGGTTTAATGCTCCAAGG-3’) is provided in the Supplementary Methods. All PCR reactions were performed using iTaq Universal SYBR Supermix (Bio-Rad, USA) in a final reaction volume of 20 µL, containing 2 µL of genomic DNA (gDNA) extracted from either a vaginal swab (as previously described) or a bacterial colony using a Chelex-based extraction method(26), 0.5 µL each of forward and reverse primers (10 µM), and molecular-grade water. Thermal cycling conditions consisted of an initial denaturation at 95 °C for 5 min, followed by 40 cycles of denaturation at 95 °C for 5 s, annealing at 65 °C for 30 s, and extension at 72 °C for 30 s. Melt curve analysis was subsequently performed from 60 to 95 °C. All samples were analysed in duplicate, and each PCR run included a positive control containing CUL40 or CUL09 gDNA and a no-template negative control. Samples were classified as positive when comparable amplification occurred in both duplicate reactions, amplicon size was confirmed by melt peak analysis and gel electrophoresis and positive and negative controls performed as expected. Species-specific primers were obtained from previously published studies: L. crispatus (forward: 5′-TCTTGACATCTAGTGCCATTTGT-3′; reverse: 5′-TGCACCACCTGTCTTAGC-3′)(27) and L. gasseri (forward: 5′-TCAAGAGCTGTTAAGGCTGT-3′; reverse: 5′-CTATCGCTTCAAGTGCTTTC-3′)(28).

### Daily Well-being diaries

Participants completed a paper-based daily diary recording the presence or absence of symptoms across four domains: vaginal (*itchiness, burning sensation, odorous discharge* and *vaginal cream/pessary use*); gastrointestinal (*abdominal pain, bloating, and indigestion*); mental (*increased irritability, increased anxiousness, increased bad mood, and increased melancholy*); and physical (*joint/muscle ache unrelated to exercise, low energy (fatigue), and poor sleep quality*).

### Statistical Analysis

Analysis used all available data (unless otherwise stated) from the ITT population and missing values were not imputed. All statistical analyses were conducted in R (v4.4.3),unless otherwise specified, and models were fit using glmmTMB (v1.1.14)(29), simulated residuals were checked using DHARMa (v0.4.7)(30) and adjusted means determined and contrasted with emmeans (v2.0.3)(31). All models include age and body mass index (BMI). For outcomes measured at multiple time points, models included group, time point, the group-by-time point interaction and, where appropriate, the baseline value of the outcome as fixed effects as well as a participant-level random intercept to account for repeated measurements from the same participant. Given the exploratory, hypothesis-generating nature of the study, *p* values were not adjusted for multiple comparisons.

#### Metagenomic sequencing

For participants with baseline and endpoint samples, the probability of participants changing CST from baseline to endpoint was compared between the groups using a two-sided Fisher’s exact test. Group differences in alpha diversity were compared at midpoint and endpoint using a covariate-adjusted linear mixed model with baseline diversity as an additional fixed effect. Beta diversity was assessed using robust Aitchison distances (vegan v2.7-5)(32) and visualised using principal coordinates analysis (PCoA). For participants with samples at all three time points, changes in the vaginal microbiota composition over time and between groups were assessed using distance-based redundancy analysis (dbRDA) conditioned on each participant to account for large inter-individual variation. The statistical significance of the changes over time was calculated through permutation of the time point label within each participant, and whether the probiotic influenced this time effect by permuting group allocation between participants while preserving each participants microbiota trajectory. Differences in community dispersion were assessed using betadisper and permutest. All permutations were done 9,999 times. Differential abundance and prevalence were analysed using covariate-adjusted MaAsLin3(33) with TSS-normalisation and log-transformation. Results associated with the group-by-time point interaction with a significant joint (abundance and prevalence outcomes combined) p value are reported. Between-group differences in the relative abundances of *L. crispatus* and *L. gasseri* were analysed using beta-binomial generalised linear mixed models (GLMMs) with a logit link, modelling the proportion of profiled reads assigned to each species.

#### Microbial culture

Between-group differences in colony-forming units (CFU) were analysed using negative binomial GLMMs with a log link. The logarithm of the fraction of the swab sample plated was used as an offset and results are expressed as CFU per swab.

#### Species and strain detection

The likelihood of detection was compared between the groups using binomial GLMMs with a logit link.

#### Daily Wellbeing Diaries

Incidence rates were calculated separately for the whole diary period (days 1-20) and the two halves (1-10 and 11-20) using generalised linear models with log-link Poisson or negative binomial families (dependent on the symptom) and an offset for the logarithm of the number of days providing data. Incidence rate ratios (IRRs) are the adjusted incidence rate in the probiotic group divided by that in the placebo group. An IRR < 1 indicates a lower rate in the probiotic group, whereas an IRR > 1 indicates a higher rate.

## Results

The flow diagram of enrolment, allocation, follow-up, and analysis is shown in Figure 1B. A total of 50 females were contacted and 40 were enrolled in January – February 2026. The probiotic and placebo were well tolerated with no reported adverse effects and no dropouts. Compliance exceeded 99% in both arms of the study. The baseline characteristics of the participants are displayed in Table 1.

**Table 1.** Baseline characteristics.

|  | **Probiotic** | **Placebo** |
| --- | --- | --- |
|  | **n=20** | **n=20** |
| Age (years), mean (SD) | 30.00 (6.78) | 31.15 (6.79) |
| Height (metres), mean (SD) | 1.68 (0.08) | 1.67 (0.07) |
| Body mass index (kg/m<sup>2</sup>), mean (SD) | 23.92 (3.75) | 22.52 (2.77) |
| Days since bleeding, mean (SD) | 2.0 (1.05) | 1.7 (0.88) |

In total, 115 swabs were collected for analysis; 5 participants (2 from probiotic group, 3 from placebo group) were unable to provide endpoint samples due to menstruation.

### Profiling of the vaginal microbiota

Shotgun metagenomic sequencing (Figure 2A) of the samples indicated that the vaginal microbial communities were dominated by *Lactobacillus* spp., particularly *L. crispatus* and *L. iners*. The microbiota profiles were categorised into Community State Types (CSTs) and at baseline were dominated by CSTs I, III and IV that were equally distributed between the groups. In the placebo group, 41% (7/17) of participants were classified as CST I, 29% as CST III and 18% as CST IV. In the probiotic group 32% (6/19) of the participants were CST I, 32% (6/19) were CST III and 32% (6/19) were CST IV (Figure 2A) and the profiles remained broadly stable throughout the intervention period (Figure 2B).

**Figure 2.**
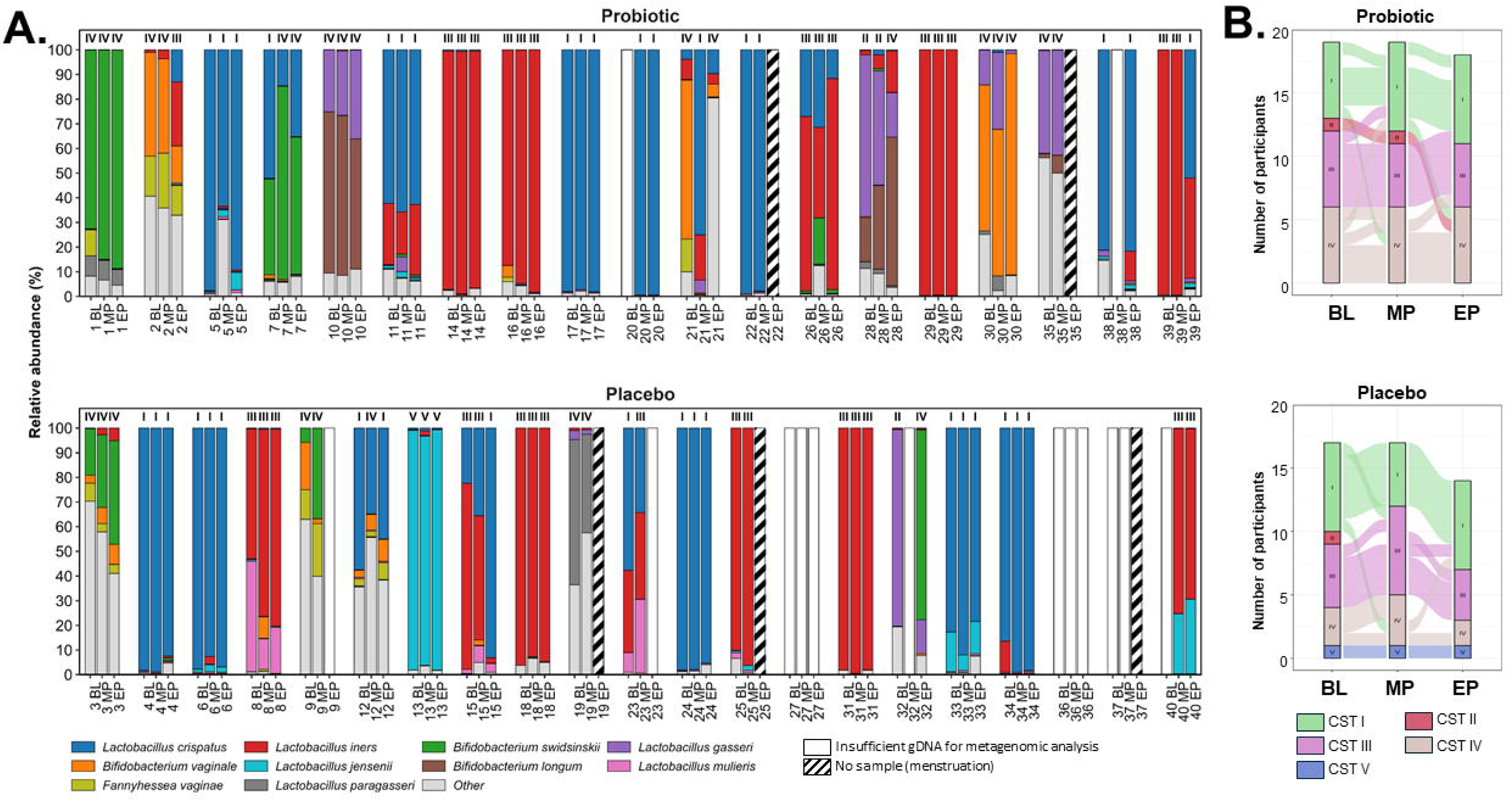
Vaginal microbiota composition and community state type (CST) transitions. **(A)** Relative abundance profiles of the 10 most abundant bacterial species and vaginal community state type (CST) classifications; **(B)** transitions in CST classification between baseline (BL), midpoint (MP) and endpoint (EP) in the placebo and probiotic groups.

Analysis of the microbiotas revealed no significant differences in alpha diversity (Figure 3A) but for beta diversity (Figure 3B) there was a significant group-by-timepoint interaction (R²=0.088; *p*=0.0177), indicating that probiotic supplementation influenced vaginal microbiota composition over time. In the probiotic group, composition differed significantly from baseline at both midpoint (R²=0.246; *p*=0.0066) and endpoint (R²=0.268; *p*=0.0090). A significant difference between baseline and endpoint was observed in the placebo group (R²=0.196; *p*=0.0322).

**Figure 3.**
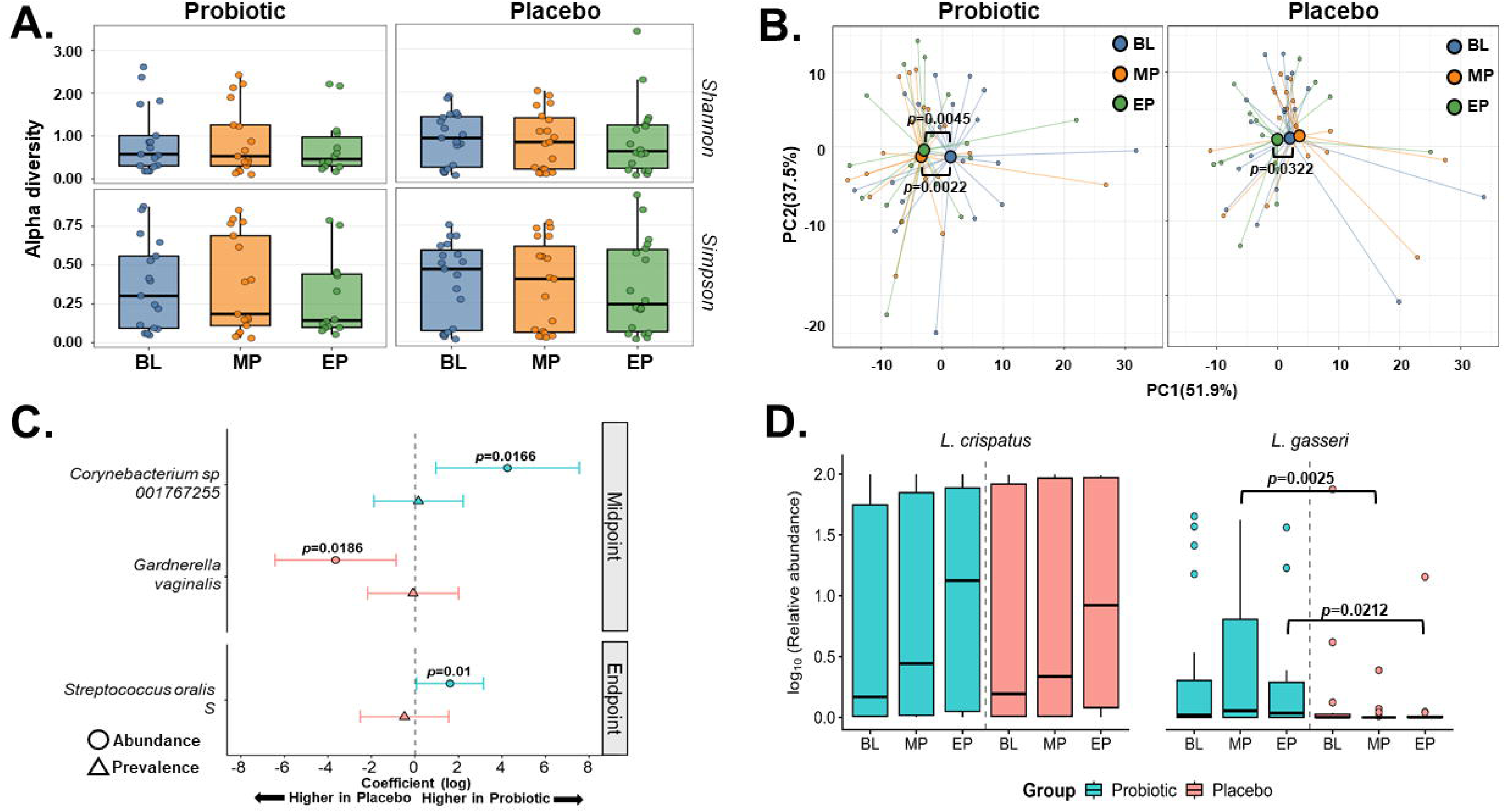
Profiling of vaginal microbiota community structure. (A) Alpha diversity (Shannon and Simpson diversity indices), **(B)** Beta diversity (principal coordinates analysis (PCoA) of robust Aitchison distances), **(C)** Changes in organisms associated with probiotic supplementation identified by differential abundance analysis, and **(D)** Relative abundance of *L. crispatus* and *L. gasseri* species. Values of *p* are as stated. BL, baseline; MP, midpoint; EP, endpoint.

Differential abundance analysis (Figure 3C) identified lower abundance of *Gardnerella vaginalis* (fold difference: 0.08; 95% CI: 0.01–0.55; *p*=0.0186) and higher abundances of *Corynebacterium* sp. 001767255 at midpoint (fold difference: 19.04; 95% CI: 1.94–186.59; *p*=0.0166) and higher abundance of *Streptococcus oralis* S at endpoint (fold difference: 3.05; 95% CI: 1.05–8.86; *p*=0.0100) in the probiotic group compared with placebo. No significant differences in species prevalence were detected.

*L. gasseri* relative abundance was similar between groups at baseline (**Figure 3D**) but was significantly higher in the probiotic group at midpoint (abundance ratio: 25.86, 95% CI: 3.14–213.09; *p*=0.0025) and endpoint (abundance ratio: 12.88; 95% CI: 1.47–113.29; *p*=0.0212). No significant between-group differences were observed for *L. crispatus*.

### Profiling of the viable vaginal microbial population

Figure 4 illustrates that there was a significant between-group difference in the viable numbers of the potentially pathogenic *Gardnerella* species at endpoint (*p*=0.0146); lower numbers were present in the probiotic group whilst the numbers in the placebo group had increased. Numbers of Enterobacteria, Enterococci and Staphylococci remained consistent during the study and there was a low background presence of the yeast, *Candida albicans*.

**Figure 4.**
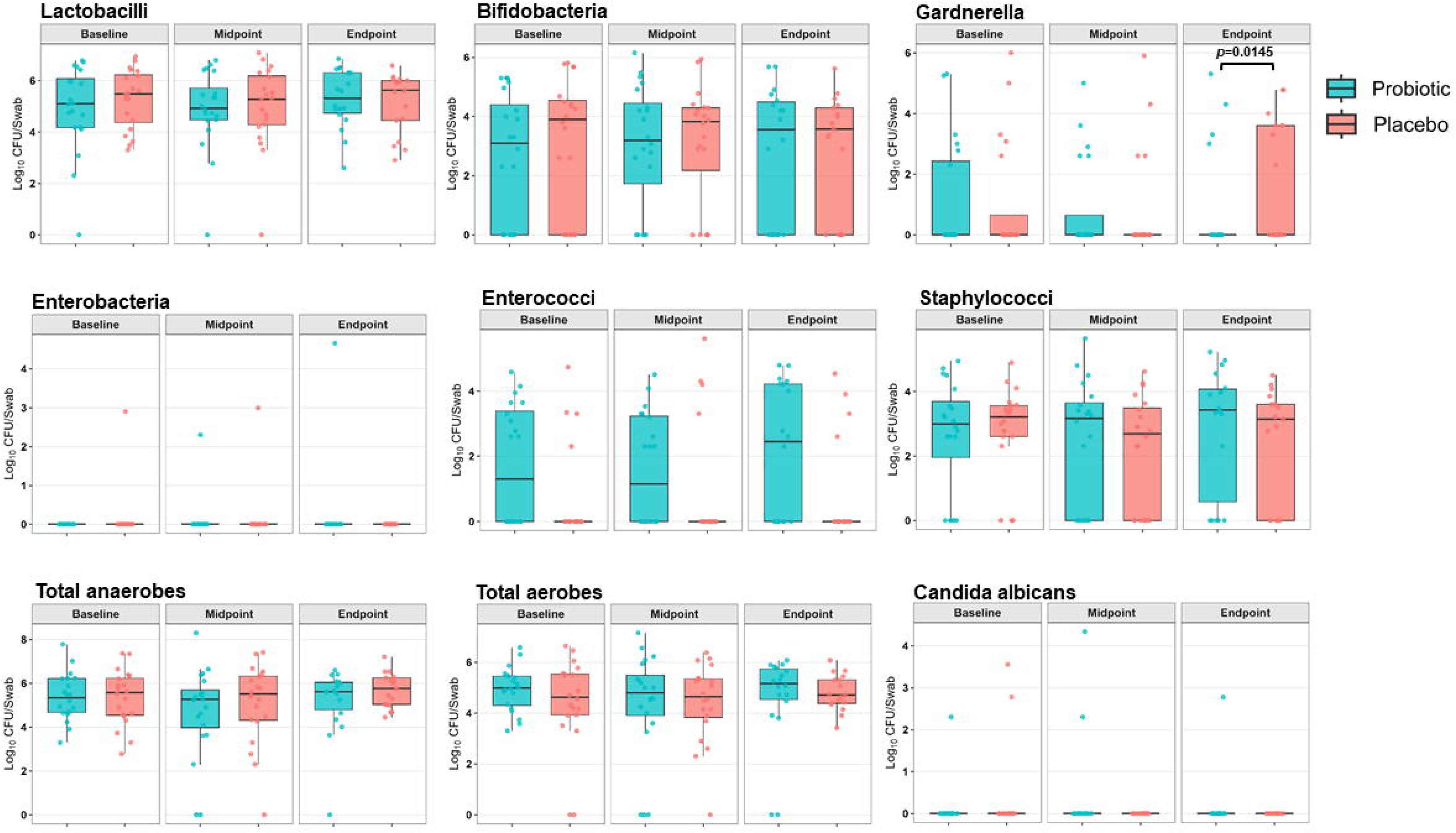
Culture-based microbiological enumeration of vaginal swabs. Values of *p* are as stated.

### Detection by qPCR of *L. crispatus* and *L. gasseri* at species level in vaginal samples

*L. crispatus* was detected in the majority of the vaginal swabs (98/115). At baseline, participant detection rates were 19/20 and 15/20 in the probiotic and placebo groups, respectively, 18/20 and 14/20 at midpoint, and 18/18 and 14/17 at endpoint (Figure 5A). No significant between-group differences were observed (Figure 5B).

**Figure 5.**
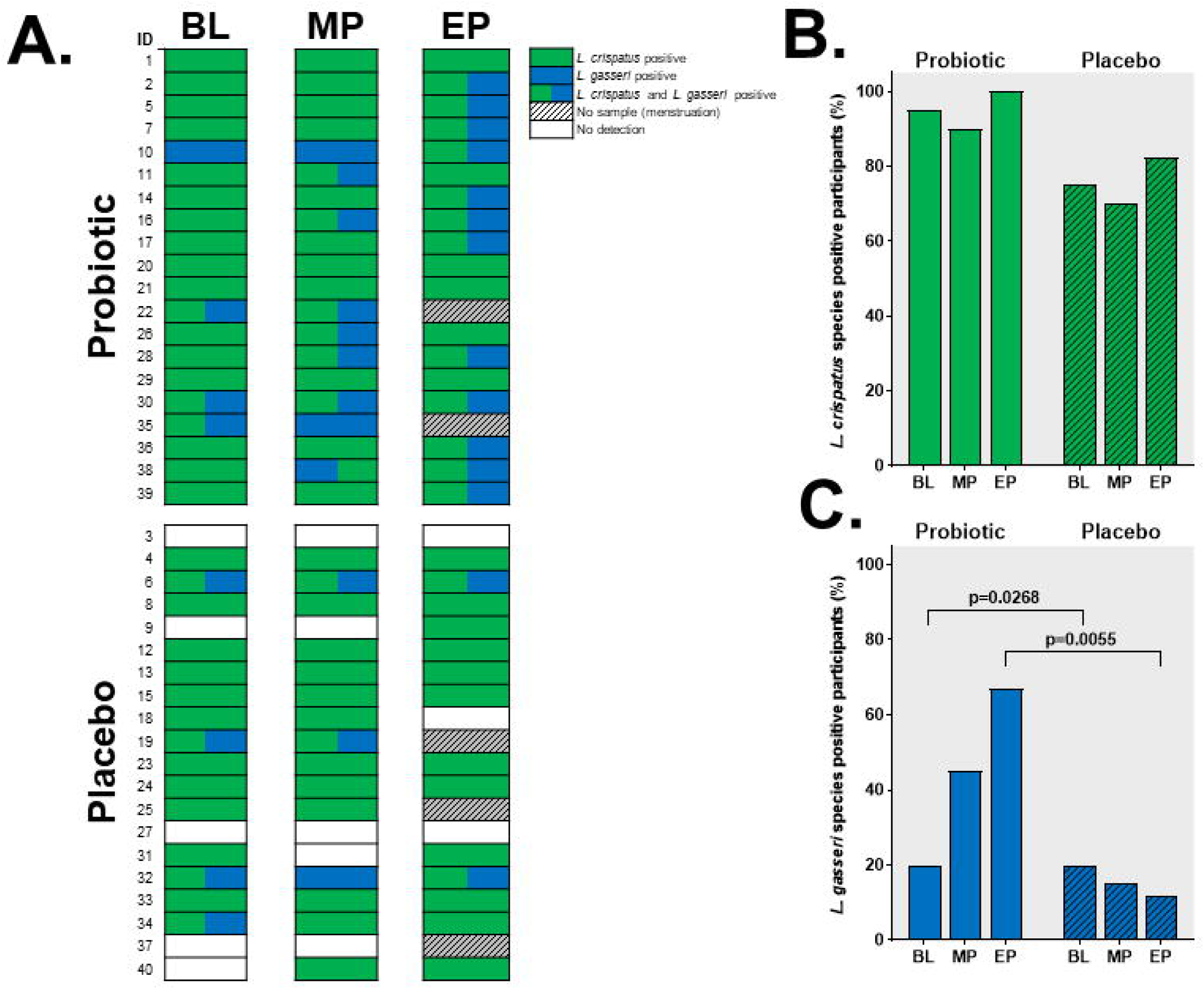
Detection and abundance of the *L. crispatus and L. gasseri* species by species-specific qPCR. **(A)** Participant-level detection of the *L. crispatus and L. gasseri* (annotated with participant ID) and the proportions of participants positive for **(B)** *L. crispatus* or **(C)** *L. gasseri*. Values of *p* are as stated.

Detection of *L. gasseri* was lower at 34/115 of the swabs. At baseline, *L. gasseri* was detected in 4/20 participants in each group, 9/20 in the probiotic group and 3/20 in the placebo group at midpoint, and 12/18 and 2/17 at endpoint (Figure 5A). Detection was significantly higher in the probiotic group at both midpoint (45% vs 15%, *p*=0.0268) and endpoint (67% vs 12%, *p*=0.0055; Figure 5C). In the probiotic group 10 participants acquired *L. gasseri* between baseline and endpoint whilst receiving the oral probiotic supplement, whilst there were no acquisitions in the placebo group.

### Detection of *L. crispatus* CUL40 and *L. gasseri* CUL09 at strain level in vaginal samples

The *L. crispatus* CUL40 strain was detected in 4/20 participants in the probiotic group and 2/20 in the placebo group at baseline indicating the presence of this strain as a component of the indigenous vaginal microbiotas of the participants (Figure 6A). At midpoint the strain was detected in 7/20 of the probiotic participants and 1/20 of the placebo with final numbers of 10/18 and 0/17 at endpoint, respectively (Figure 6A). Each of the *L. crispatus* CUL40-positive participant detections corresponded with the detection of *L. crispatus* at the species level (Figure 5). At midpoint, *L. crispatus* CUL40 was detected in 35% of the probiotic group and in 5% of the placebo group (*p*=0.0301; Figure 6B). At endpoint, CUL40 was detected in 55.6% of the participants in the probiotic group and was not detected in the placebo group (*p*=0.0050). Seven participants in the probiotic group had acquired CUL40 by endpoint. CUL40 was detected at baseline but not at endpoint in two participants in the placebo group and one in the probiotic group.

**Figure 6.**
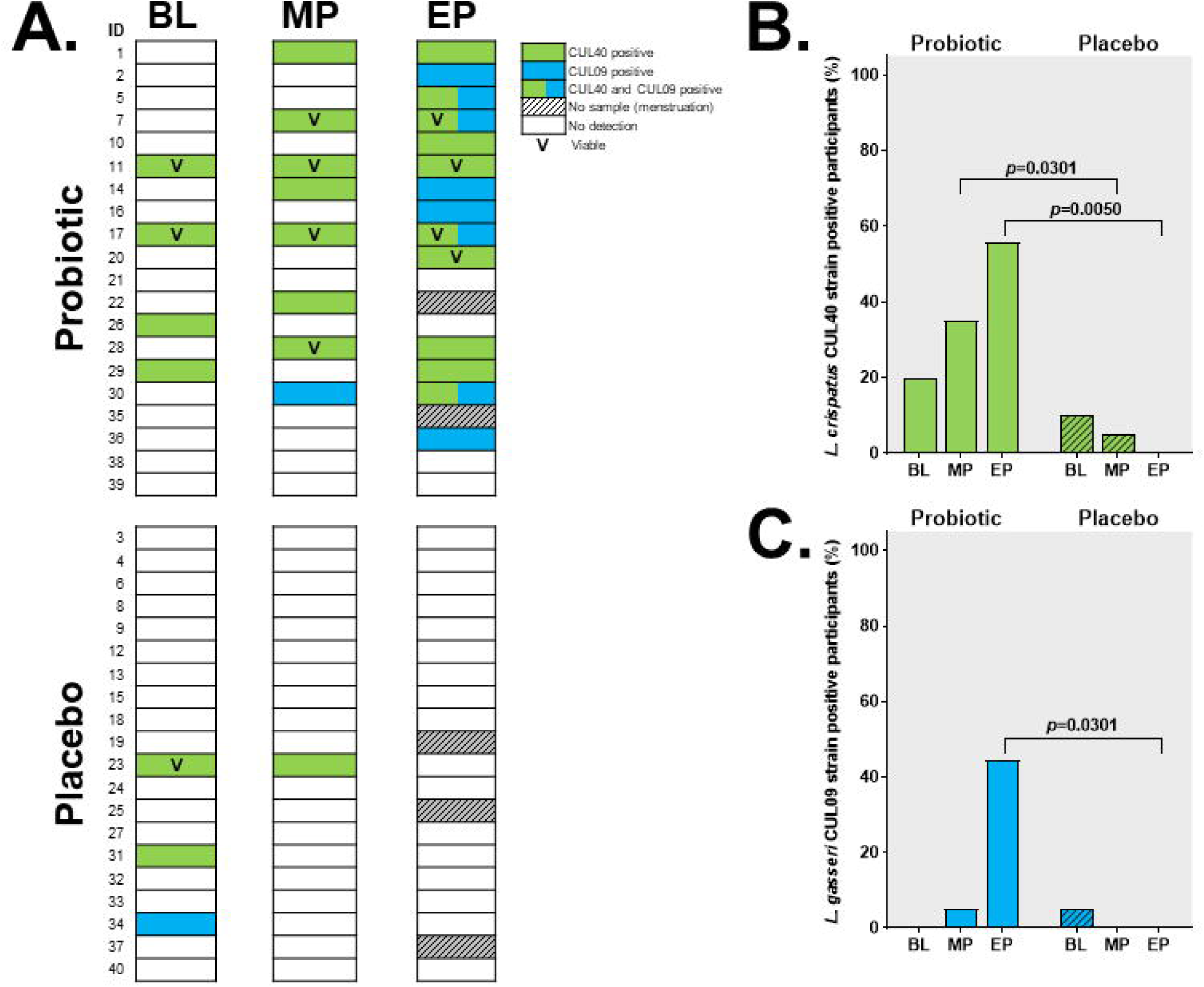
Detection and abundance of *L. crispatus* CUL40 and *L. gasseri* CUL09 by strain-specific qPCR. **(A)** Participant-level detection of the *L. crispatus* CUL40 and *L. gasseri* CUL09 strains (annotated with participant ID) and the proportions of participants positive for **(B)** *L. crispatus* CUL40 or **(C)** *L. gasseri* CUL09. Values of *p* are as stated.

The *L. gasseri* CUL09 strain was detected in one participant (in the placebo group) at baseline, and by the end of the intervention it was detected in 8/18 participants in the probiotic group and none of the placebo (Figure 6A); all strain detections were confirmed at species level (Figure 5). At endpoint, CUL09 detection was significantly higher in the probiotic group than in the placebo group (44.4% vs 0%, p=0.0301; Figure 6C). All eight CUL09-positive participants in the probiotic group had acquired CUL09 during the intervention. Three participants acquired both CUL40 and CUL09. CUL40 was detected at baseline but not at endpoint in two participants in the placebo group and one in the probiotic group. CUL09 was detected at the baseline but not at the endpoint for one participant in the placebo group.

The MRS agar plates for the samples testing positive for either CUL40 or CUL09 were screened for the presence of viable colonies of the strains using strain-specific qPCR. Viability was confirmed for 46% (11/24, Figure 6A) of the CUL40-positive participants including midpoint/endpoint samples from three participants in the probiotic group who had acquired CUL40. Viable colonies of CUL09 were not detected.

### Daily Well-being Diaries

Symptoms reported during the intervention period are presented in Figure 7. During the second half of the intervention (days 11–20), there were fewer reported symptoms in the probiotic group than in the placebo group. Vaginal itchiness (Figure 7A) and bloating (Figure 7B), were completely absent in the probiotic group during this period but were present in the placebo. Reports of vaginal cream or pessary use were infrequent.

**Figure 7.**
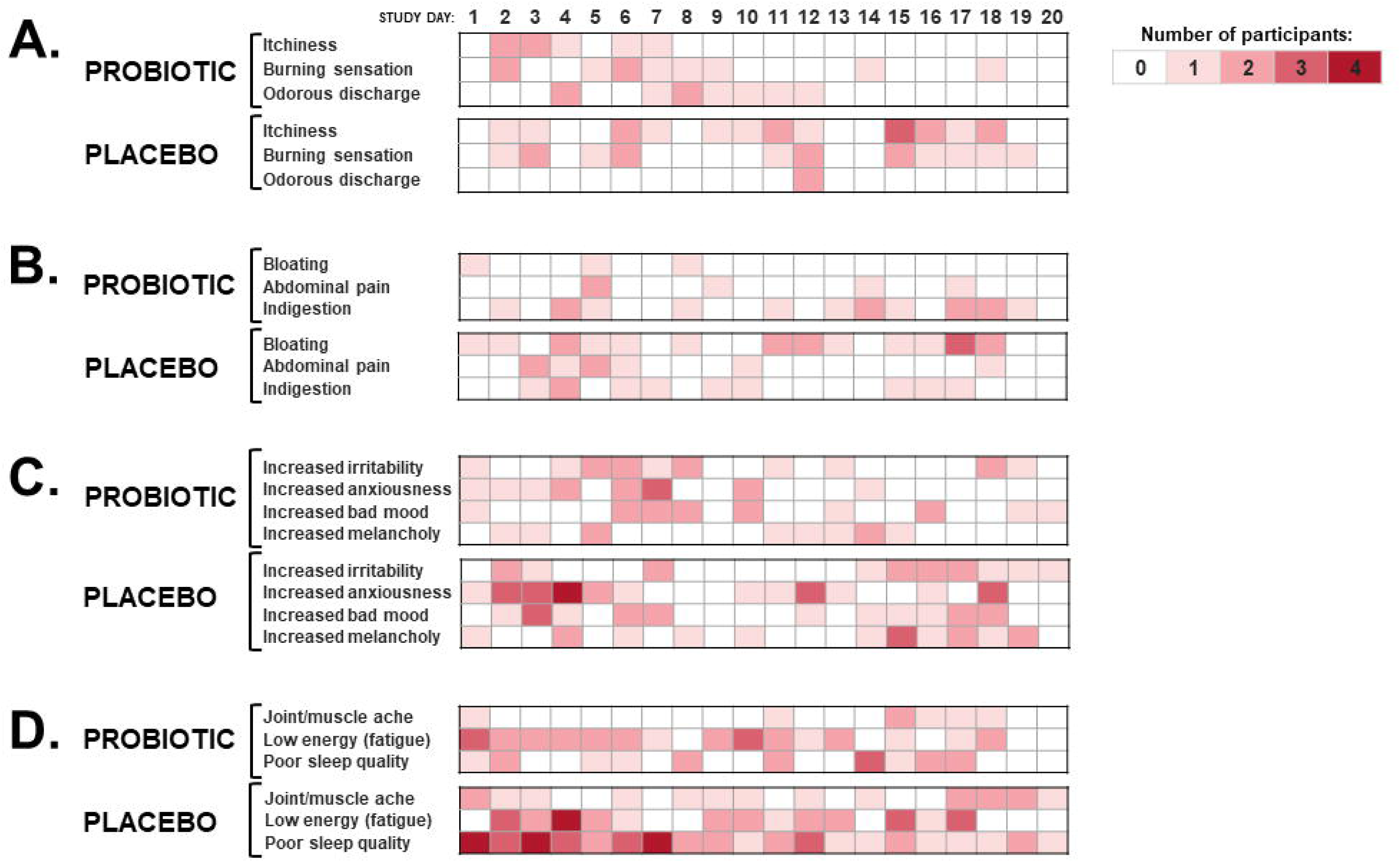
Daily symptom reporting in the probiotic and placebo groups. Heatmaps showing daily symptom reports in the probiotic and placebo groups grouped into (A) vaginal, (B) gastrointestinal, (C) mental and (D) physical domains.

Incidence Rate Ratios (IRRs, probiotic vs placebo) of symptoms during the second half of the study are shown in Table 2 along with IRRs for the first half and the full intervention period.

**Table 2.**
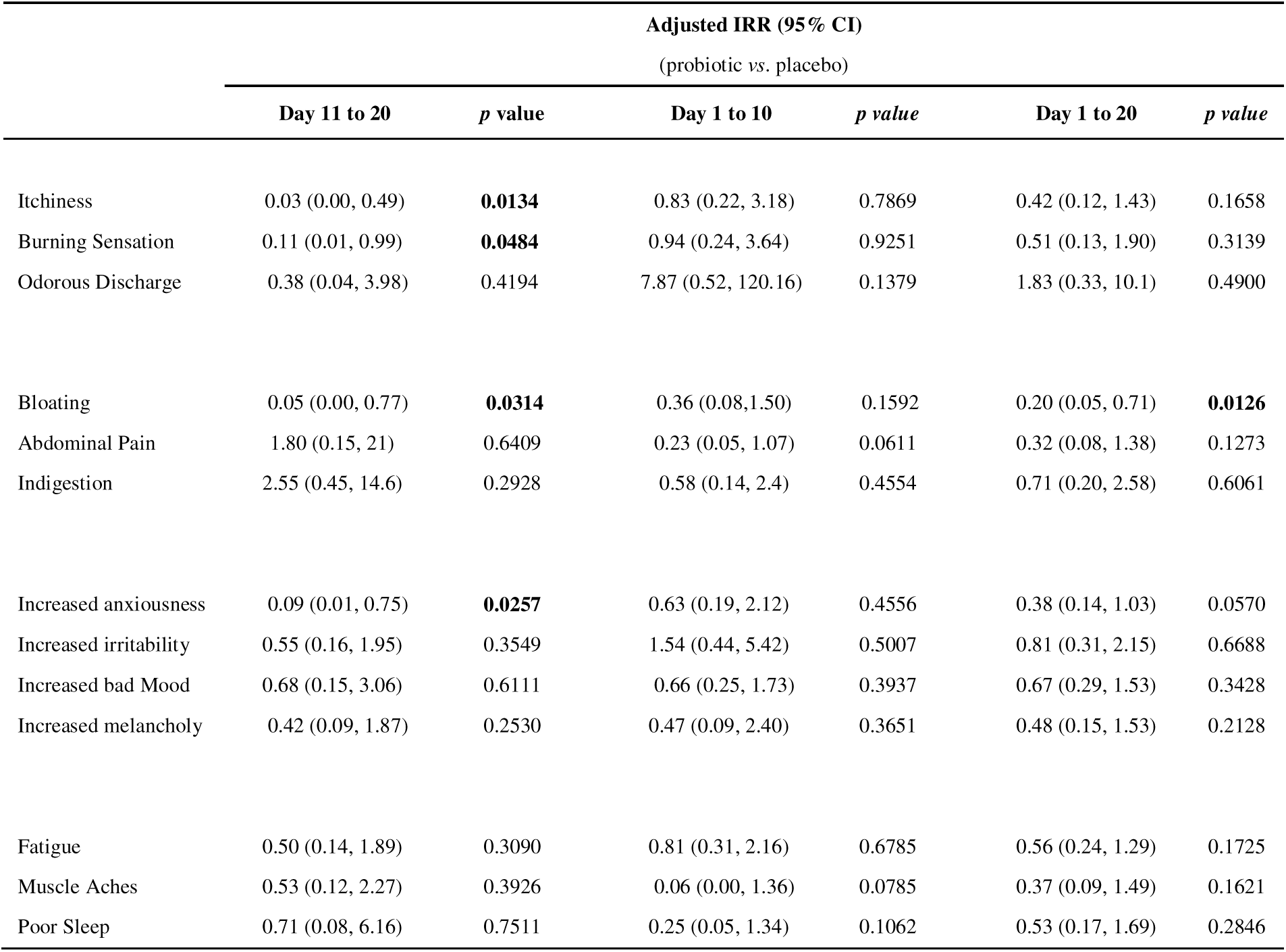
Adjusted Incidence Rate Ratios for symptoms.

|  | Adjusted IRR (95% CI)<br>(probiotic vs. placebo) |  |  |  |  |  |
| --- | --- | --- | --- | --- | --- | --- |
|  | Day 11 to 20 | *p* value | Day 1 to 10 | *p* value | Day 1 to 20 | *p* value |
| Itchiness | 0.03 (0.00, 0.49) | **0.0134** | 0.83 (0.22, 3.18) | 0.7869 | 0.42 (0.12, 1.43) | 0.1658 |
| Burning Sensation | 0.11 (0.01, 0.99) | **0.0484** | 0.94 (0.24, 3.64) | 0.9251 | 0.51 (0.13, 1.90) | 0.3139 |
| Odorous Discharge | 0.38 (0.04, 3.98) | 0.4194 | 7.87 (0.52, 120.16) | 0.1379 | 1.83 (0.33, 10.1) | 0.4900 |
| Bloating | 0.05 (0.00, 0.77) | **0.0314** | 0.36 (0.08, 1.50) | 0.1592 | 0.20 (0.05, 0.71) | **0.0126** |
| Abdominal Pain | 1.80 (0.15, 21) | 0.6409 | 0.23 (0.05, 1.07) | 0.0611 | 0.32 (0.08, 1.38) | 0.1273 |
| Indigestion | 2.55 (0.45, 14.6) | 0.2928 | 0.58 (0.14, 2.4) | 0.4554 | 0.71 (0.20, 2.58) | 0.6061 |
| Increased anxiousness | 0.09 (0.01, 0.75) | **0.0257** | 0.63 (0.19, 2.12) | 0.4556 | 0.38 (0.14, 1.03) | 0.0570 |
| Increased irritability | 0.55 (0.16, 1.95) | 0.3549 | 1.54 (0.44, 5.42) | 0.5007 | 0.81 (0.31, 2.15) | 0.6688 |
| Increased bad Mood | 0.68 (0.15, 3.06) | 0.6111 | 0.66 (0.25, 1.73) | 0.3937 | 0.67 (0.29, 1.53) | 0.3428 |
| Increased melancholy | 0.42 (0.09, 1.87) | 0.2530 | 0.47 (0.09, 2.40) | 0.3651 | 0.48 (0.15, 1.53) | 0.2128 |
| Fatigue | 0.50 (0.14, 1.89) | 0.3090 | 0.81 (0.31, 2.16) | 0.6785 | 0.56 (0.24, 1.29) | 0.1725 |
| Muscle Aches | 0.53 (0.12, 2.27) | 0.3926 | 0.06 (0.00, 1.36) | 0.0785 | 0.37 (0.09, 1.49) | 0.1621 |
| Poor Sleep | 0.71 (0.08, 6.16) | 0.7511 | 0.25 (0.05, 1.34) | 0.1062 | 0.53 (0.17, 1.69) | 0.2846 |

During days 11–20, significantly less vaginal itchiness (adjIRR: 0.03, *p*=0.0134) and burning sensation (adjIRR: 0.11, *p*=0.0484) were reported by the probiotic group compared to the placebo, corresponding to reductions of 97% and 89% respectively. Bloating was 95% lower in the probiotic group during this period (adjIRR: 0.05, *p*=0.0314), with a significant reduction also observed for the full 20-day intervention period (adjIRR: 0.20, *p*=0.0126). In addition, anxiousness was 91% lower in the probiotic group than in the placebo group during days 11–20 (adjIRR: 0.09, *p*=0.0257).

## Discussion

At the end of the 20-day double-blind, placebo-controlled study with the Lab4V probiotic, both the vaginal strains (*Lactobacillus crispatus* CUL40 and *Lactobacillus gasseri* CUL09) were detected in the vaginal samples provided by the participants in the probiotic group but neither was detected in the placebo group. There were changes in the vaginal microbiota composition in the probiotic group including a reduced presence of *Gardnerella* spp. and, in this group, there were fewer reports of vaginal itchiness/burning sensation.

CSTs remained broadly stable during the course of the study and there were no significant changes in the alpha diversity outcomes. Analysis of the beta-diversity identified a significant group-by-timepoint interaction, indicating changes in microbial community composition. Culture-based analysis identified significantly lower *Gardnerella* spp. counts in the probiotic group at endpoint and *L. gasseri* has been shown to inhibit *Gardnerella vaginalis*(34). Recording vaginal pH would have provided additional context for interpreting these findings.

The strain-specific PCR analysis at baseline identified a background presence (17.5% (7/40) of participants) of one or both the Lab4V vaginal strains suggesting a low level of prevalence within the indigenous vaginal microbiota of the general population. By the study endpoint, detection rates in the probiotic group had increased to 55.6% (10/18) for *L. crispatus* CUL40 and 44.4% (8/18) for *L. gasseri* CUL09, whereas neither strain was detected in the placebo group. Seven participants acquired CUL40 and eight acquired CUL09 by endpoint, with three participants acquiring both strains. Acquisition of the targeted strains was independent of baseline CSTs, indicating that detection of the supplemented strains was not restricted to a particular baseline vaginal community composition. In a study by Perez *et al*. (2025), oral administration of a different strain (*L. gasseri* CECT 30648) resulted in detection in the vaginal microbiota of 55.9% of probiotic recipients whilst the co-administered *L. crispatus* strain was not detected(35). Other studies have failed to demonstrate vaginal detection of orally supplemented lactobacilli(36), which suggests strain specificity.

Here, strain-specific PCR was combined with culture-based colony screening to confirm the viability of the strains translocated to the vagina as such evidence is very limited(37–40). Using this approach, it was possible to detect viable *L. crispatus* CUL40 but that could not be achieved for *L. gasseri* CUL09. This could be a limitation of the methodology and further work is needed.

From days 11 – 20 the incidence of both vaginal itchiness and burning sensation in the probiotic group was minimal but that was not the case in the placebo group where the rates were consistent throughout the study. Beneficial effects of orally administered probiotics on vaginal health have been seen with *Lacticaseibacillus rhamnosus* CA15(13) that resulted in modulation of the vaginal microbiota composition and improvements in the symptoms of vaginal dysbiosis. A meta-analysis of 10 randomised trials involving 1,234 women reported a 45% reduction in the incidence of bacterial vaginosis recurrence following probiotic supplementation(41) and a separate meta-analysis reported improved bacterial vaginosis cure rates in trials involving oral probiotics, although substantial heterogeneity was observed(42). Conversely, Lyra et al. (2023) found no significant changes in vaginal microbiota composition following oral supplementation in healthy women with *L. acidophilus* La-14 and *L. rhamnosus* HN001(36).

There were reductions in the occurrence rates of bloating and of anxiousness in the participants receiving the probiotics, suggesting that the potential benefits gained from probiotic intake may extend beyond vaginal health. These findings are consistent with previous studies with the Lab4 probiotic consortium that have indicated improvements in indicators of intestinal health and anxiety in both IBS sufferers and healthy adults(19, 43). Meta-analyses of probiotic studies have reported improvements for bloating and anxiety which are formulation dependent although the certainty of evidence is limited(44–47).

In conclusion, 20 days of oral supplementation, between menstruation cycles, with a six-strain probiotic formulation resulted in increased vaginal detection of the orally supplemented strains of *L. crispatus* CUL40 and *L. gasseri* CUL09, changes in vaginal microbial composition, reduced presence of *Gardnerella* spp. and improvements in the levels of vaginal discomfort. The recovery of viable CUL40-positive bacteria provides further evidence supporting the potential for orally administered lactobacilli to translocate into the vaginal ecosystem. The outcomes from this pilot study provide a strong basis for future full-scale trials.

## Acknowledgements

This study was funded by Cultech Ltd, which also supplied the study intervention. The funder contributed to study design, data analysis and interpretation, manuscript preparation, and the decision to submit the manuscript for publication.

## Authors’ contributions

Conceptualization: DAJ, DRM, NC, SP. Methodology: DAJ, NC Investigation: EH, DAJ, NC, DRM Formal Analysis: EH, DAJ, NC, DRM, JMDS, ES, EG, DW. Data curation: MD. Validation: DAJ, DRM, NC, SP Writing-original draft: DAJ, DRM, NC, SP, EH Writing-review and editing: EH, DAJ, DRM, NC, JMDS, MD, DW, ES, EG, SP Supervision: DRM, SP. All authors have approved the final version of the manuscript.

## Ethics approval and consent to participate

The studies involving humans were approved by the Clarenta Ethics committee, (Reference: No.001/27.11.2025r). The study was registered with clinicaltrials.gov (ID: NCT07364383, 06/01/2026). The study was conducted in accordance with the Declaration of Helsinki. All participants provided their written informed consent to participate in this study.

## Availability of data and materials

The sequencing data presented in this study can be found in online repositories. The names of the repository/repositories and accession number(s) can be found below: https://www.ebi.ac.uk/ena, PRJEB128808.

