## Supplementary Table 1 for "A Pilot Study Investigating the Impact of Oral Probiotic Supplementation on the Vaginal Microbiota and Aspects of Well-being"

**SUPPLEMENTARY INFORMATION**

**Supplementary Table S1. Selective microbial culture conditions.**

| **Culture** | **Selective medium** | **Growth conditions** |
| --- | --- | --- |
| Lactobacilli | De Man, Rogosa and Sharpe (MRS) agar (CM0361) | Anaerobic*, 37 °C for 72 h |
| Bifidobacteria | MRS-X agar (SR0051C) | Anaerobic*, 37 °C for 72 h |
| Enterobacteria | MacConkey No. 3 agar (CM0115B) | Aerobic, 37 °C for 48 h |
| Enterococci | Slanetz & Bartley (CM0377B) | Aerobic, 37 °C for 48 h |
| Streptococcus | KF Streptococcus (CM0701B) | Aerobic, 37 °C for 48 h |
| Staphylocci | Baird Parker agar + egg yolk tellurite emulsion supplement (CM0275B, SR0054) | Aerobic, 37 °C for 48 h |
| Gardnerella | Gardnerella selective agar (MED1248) | Anaerobic*, 37 °C for 48 h |
| Candida | Candida CHROMID (Biomerrieux, 43631) | Aerobic, 37 °C for 24 h |
| Total Aerobes | Colombia + 5% HBA (Biomerieux, 43050) | Aerobic, 37 °C for 48 h |
| Total Anaerobes | Horse Blood Agar | Anaerobic*, 37 °C for 48 h |

*10% carbon dioxide, 5% hydrogen and 85% nitrogen.
