## Supplementary Methods for "A Pilot Study Investigating the Impact of Oral Probiotic Supplementation on the Vaginal Microbiota and Aspects of Well-being"

**Supplementary Methods: Design of strain-specific qPCR assays for *Lactobacillus gasseri* CUL09 and *Lactobacillus crispatus* CUL40**

***Design of strain-specific primers***

Genomic DNA was extracted from overnight cultures of *Lactobacillus gasseri* CUL09 and *Lactobacillus crispatus* CUL40 using the QIAamp® PowerFaecal Pro DNA Kit (Qiagen, Germany). Long-read sequencing was performed using the PacBio Revio platform (Novogene Cambridge, UK), and PacBio SMRT Link software to generate a high-quality long-read genome assemblies.

Candidate target-specific loci were identified using regions present in the target genome but absent from the comparator background detected by whole-genome alignment with NUCmer^1^. Candidate primer sequences targeting candidate target-specific loci were designed using Primer3^2^. The predicted amplicon sequences are displayed in **Table S1**. The specificity of candidate primer pairs was assessed using BLASTn against the NCBI nucleotide collection (accessed November 2025) to ensure the absence of off-target amplicons. Both primer pairs targeted single copy genes encoding hypothetical proteins. The targeted coding sequences were located at positions 566,260–566,673 in CUL40 and 1,461,864–1,464,935 in CUL09.

**Table S1. Predicted amplicon sequences for CUL40 and CUL09 primers**

| **Primer** | **Amplicon sequence (5’-3’) with primer binding sites in grey** | **Amplicon size (bp)** |
| --- | --- | --- |
| CUL40 | TGGACTGATTCCCCCTTTGACATGCACACTTATAAAAAAGGCGATGAAGCTCGTTGCATTGGCGCAGATGACGATTACTACGTTGTTGATGACCTCCCA | 99 |
| CUL09 | TATGGTCGGTTAACACGGGTCAAAGCTTGCAATAAGTTATGCGCTTTGATCATTCGTCCAAGGTATAATTTTTTTAGCCTTGGAGCATTAAACCCCGT | 98 |
